# Graph Attention Site Prediction (GrASP): Identifying Druggable Binding Sites Using Graph Neural Networks with Attention

**DOI:** 10.1101/2023.07.25.550565

**Authors:** Zachary Smith, Michael Strobel, Bodhi P. Vani, Pratyush Tiwary

## Abstract

Identifying and discovering druggable protein binding sites is an important early step in computer-aided drug discovery but remains a difficult task where most campaigns rely on *a priori* knowledge of binding sites from experiments. Here we present a binding site prediction method called Graph Attention Site Prediction (GrASP) and re-evaluate assumptions in nearly every step in the site prediction workflow from dataset preparation to model evaluation. GrASP is able to achieve state-of-the-art performance at recovering binding sites in PDB structures while maintaining a high degree of precision which will minimize wasted computation in downstream tasks such as docking and free energy perturbation.

## INTRODUCTION

A critical early step in computer-aided drug discovery is identifying druggable binding sites or those that can bind ligands likely to alter activity. Virtual screening of ligands with docking methods is often done for a specific binding site which requires *a priori* knowledge of where ligands are likely to bind.^1–4^ Recently, modern structure prediction methods such as AlphaFold2^5,6^ and RoseTTAFold^7^ have greatly expanded the number of predicted structures for the human proteome^8^ while enhanced sampling methods for molecular dynamics have revealed conformations with cryptic pockets inaccessible in the protein’s crystal structure.^9–11^ The combination of advances in these two areas has led to a deluge of protein conformations that have not been probed for binding sites in experiments. For drug discovery to keep pace with structure discovery, accurate high-throughput binding site identification methods must be developed.

Initially, binding site prediction methods used human-designed representations of proteins based on geometry,^12–18^ sequence conservation,^19,20^ interactions with probe molecules,^21,22^ or a combination of these features.^2,23^ Recent methods, however, have leveraged machine learning combined with binding-site databases^24,25^ to learn how to predict binding sites.^26–33^ Despite the existence of large databases and modern machine learning architectures, one of the most popular and successful methods in this area is P2Rank, a random forest classifier trained on 251 protein structures.^27^ It is striking that this model is able to outperform a Convolutional Neural Network (CNN) trained on thousands of structures.^26^ The reason behind P2Rank’s success might be the use of better representations such as an accessible surface area mesh with a rotationally invariant model or the use of a smaller but more carefully curated dataset. One more recently developed class of machine learning architectures that employs a natural representation for molecules are Graph Neural Networks (GNNs)^34,35^ which represent inputs as graphs and pass messages between connected nodes. GNNs have been shown to excel at closely related tasks such as binding affinity prediction,^36,37^ docking,^38^ predicting which sites will open mid-simulation,^39^ predicting the type of molecule that binds to a known site,^40^ and even predicting protein-protein interactions.^41^ Like P2Rank, GNNs also have rotational invariance guaranteeing the orientation of an input molecule does not affect the internal representation. With this motivation, we have developed a GNN-based method called Graph Attention Site Prediction (GrASP). GrASP is designed with the representational advantages of P2Rank in mind and performs a rotationally invariant featurization of solvent-accessible atoms. As a deeper model, GrASP requires a larger dataset for training, and to achieve this goal we have created a new publicly available version of the sc-PDB database containing 26,196 binding sites across 16,889 protein structures. GrASP is able to recover a higher number of ground truth binding sites when evaluated on P2Rank’s test sets but has the important advantage that over 70% of its output binding sites correspond to real binding sites whereas under 30% of P2Rank sites correspond to real sites.

## METHODS

In this section, we introduce Graph Neural Networks and show each step of the site prediction pipeline including dataset creation, protein representation, and the model architecture.

### Graph Neural Networks (GNNs)

For the sake of better motivating the architecture underlying GrASP, we start with a brief pedagogical overview. Graph Neural Networks (GNNs) are a family of architectures that operate on a graph structure to represent the features of individual nodes and the relational structure between them. In this work, we represent proteins as graphs in which nodes represent heavy atoms, and edges are drawn between all pairs of atoms within 5 Å of each other. Node features include both atomic features such as formal charge and residue features such as residue name. Edges also have features of inverse distance and bond order. A full list of features can be found in the Supporting Information (SI). GNNs featurize nodes using message-passing layers which perform the following three operations:

1. Message: Neighboring nodes send information to one another about their current state.
2. Aggregate: Each node collects the messages from its neighbors and aggregates them by applying an aggregation function.
3. Update: Each node incorporates the aggregated information with its own representation to generate a new latent representation of itself.

This process can be formalized as the following:^42^

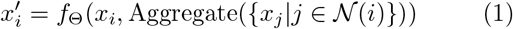

Here *x*_*i*_ is the current representation of node *i*, 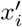 is the updated representation of node *i*, 𝒩 (*i*) denotes the set of neighbors connected to node *i*, and *f*_Θ_ denotes a parameterized update function.

This process can be repeated with multiple GNN layers for a node’s representation to incorporate information from a larger region of the graph. Since each message includes information about a node’s immediate neighbors, each GNN layer allows the node to access information influenced by nodes one hop further than the previous layer.^43^ This can be seen in Fig. 1 where the inference node’s hidden representation would include information about *k*-hop neighbors after passing through *k* GNN layers. These repeated GNN layers are commonly used within an encoder-processor-decoder framework implemented through multilayer perceptrons (MLPs) before and after a set of GNN layers.^44^

**FIG. 1:**
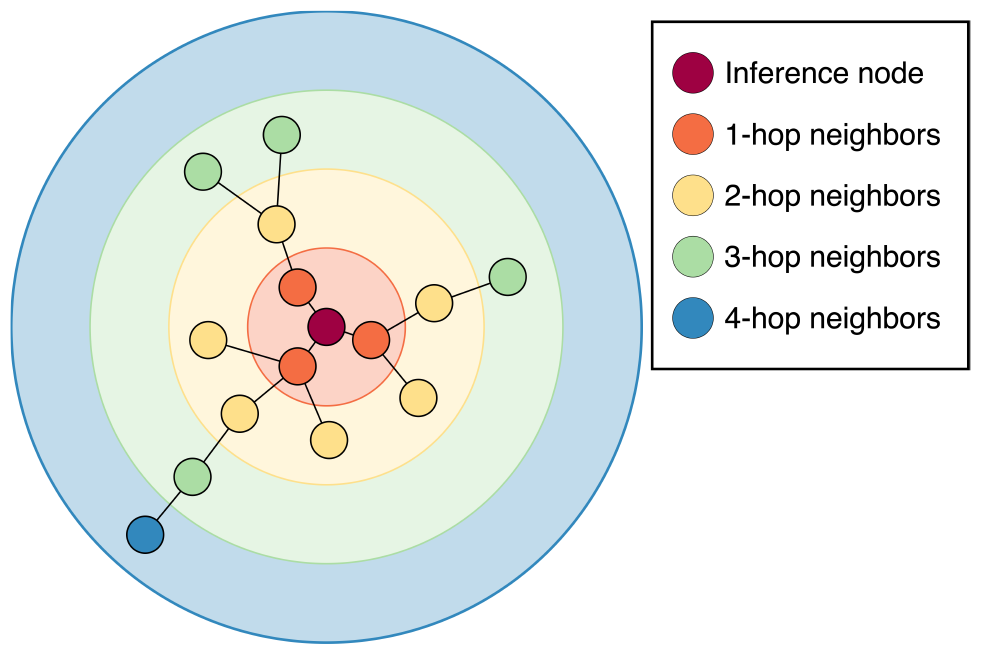
*k*-hop neighborhoods for a given inference node in the input graph. The *k*-th GNN layer representation is affected by neighbors up to *k* hops away.

Repeated aggregation comes at the cost of oversmoothing, a phenomenon where deeper GNNs cause node representations to become increasingly similar.^45^ A number of methods have been developed to encourage diverse latent representations and allow for deeper GNN architectures. Three of these are used in this work: ResNet skip connections,^46^ jumping knowledge skip connections,^47^ and Noisy Nodes.^45^ Both ResNet and jumping knowledge skip connections preserve information from earlier GNN layers (equivalently *k*-hop neighborhoods) by combining their latent representations with those of later layers. ResNet skip connections do so locally by adding the input and output of each GNN layer while jumping knowledge skip connections feed the latent representations of multiple GNN layers into the decoder. In contrast, Noisy Nodes is a regularization procedure where noise is added to the input features, and an additional decoder head that attempts to reconstruct the de-noised inputs is added after the processor layers, forcing the intermediate processor layer’s latent representations to maintain enough diversity to reconstruct inputs.

### Graph Attention Networks (GAT)

Graph attention networks (GAT) are GNNs that use attention to learn weights for each neighbor and perform a weighted average aggregation.^48^ A GAT layer is shown in Eq. 2 where Θ is a linear layer and *α*_*i,j*_ represents the attention coefficient for messa from node *j* to node *i*.

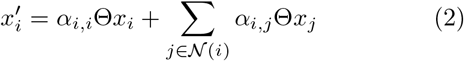

We use the attention function from GATv2 which calculates weights with the softmax of an MLP over a concatenation of both node and edge features.^42^ This function is shown in Eq. 3 where || represents concatenation, *e*_*i,j*_ are edge features and the linear layers Θ and *a*^⊺^ form the MLP.

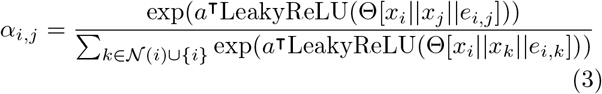

### Graph Attention Site Prediction (GrASP)

GrASP is a GAT-based model for binding site prediction. GrASP first employs the GAT model to perform semantic segmentation on all protein surface atoms, scoring which atoms are likely part of a binding site. These atomic scores are then aggregated into binding sites using average linkage clustering^49^ and ranked as a function of their constituent atoms’ scores. This overall workflow performs an instance segmentation task (binding site prediction) by postprocessing the semantic segmentation predictions (atomic binding scores).

#### Preprocessing

The first issue we address is the definition of a binding site, for which there is no consensus definition in the literature. Definitions range from atoms within 2.5 Å^50^ of the ligand to residues within 6.5 Å^24^ and choose to include different combinations of empty space, surface atoms (or surface meshes), and buried atoms. This wide range of representations has two implications. The first implication is that we can not perform an unbiased comparison with metrics based on a specific definition because we would artificially skew success rates toward methods trained with a similar definition. For example, one metric we can not use is the volume overlap between the “true” and predicted binding sites. We focus on a metric that directly compares predictions to the ligand instead of a prescribed area around it: the distance from the predicted site center to any ligand-heavy atom. This is not the only metric that fits this criterion but we choose to use it for fair comparison because P2Rank was also tuned using this metric. The second implication of not having a consensus binding site definition is that we can tune the definition used during training to maximize the model’s performance on our chosen metrics. Since these metrics do not rely on the site definition, we can tune this hyperparameter without affecting the evaluation of other methods. To achieve this goal, we assign a continuous target score to each surface atom using a sigmoid function on the distance between the ligand and protein atom. This representational choice, for which we provide details in the SI, makes it so that GrASP is penalized more for incorrectly characterizing atoms near ligands instead of treating all atoms within a cutoff distance as the same.

The second issue we address is defining the protein graph. We do this using the same inductive bias as the binding site definition: only surface atoms can be considered binding sites. This means that we will only score surface atoms but we wish to characterize the local chemical environment of these atoms using their neighbors. We construct a near-surface graph consisting of both surface atoms, defined using solvent-accessible surface area, and buried atoms within 5 Å of surface atoms. In other words, we use the induced sub-graph consisting of the surface atoms’ one-hop neighborhood. More precise details about the implementation of this representation are available in the SI. This representational choice gives GrASP the inductive bias that only surface atoms are accessible and allows it to learn druggability without first learning which atoms a ligand can reach.

#### Architecture

It has been shown that there is no best aggregator for graphs with continuous features.^51^ This has led to the development of GNNs using multiple aggregators. This multi-aggregation strategy is the inspiration for the GrASP block shown in Fig. 2A. This block consists of a GAT layer with four attention heads that pass both summed and averaged messages through a linear layer, an InstanceNorm,^52^ a residual skip connection,^46^ and an Elu activation.^53^ The linear layer after the multi-aggregation allows the model to decide how much weight to give the sum and mean for each feature.

**FIG. 2:**
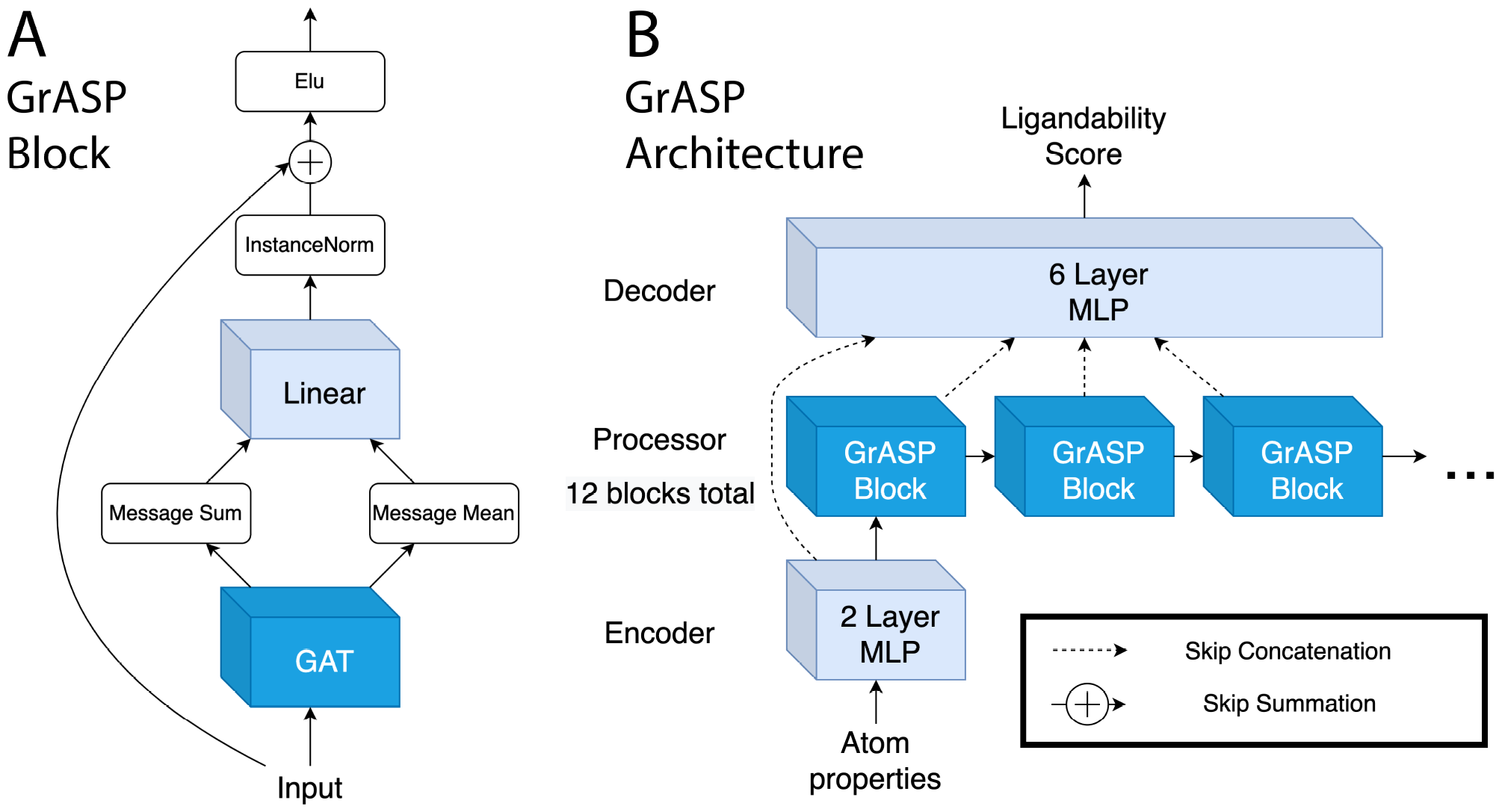
Diagram of the GrASP model. A) The GrASP blocks used to represent each atom’s local chemical environment. B) The full architecture combining GrASP blocks in an encoder-processor-decoder framework. Layers that do not consider neighbors are light blue while layers that consider neighbors are blue.

These GrASP blocks are combined with an MLP encoder and MLP decoder to make the full GrASP model shown in Fig. 2B. The output of each hybrid block is concatenated using jumping knowledge skip connections^47^ as an input for the decoder. During training, GrASP also receives inputs with Gaussian noise added and uses a second Noisy Nodes^45^ head to reconstruct denoised inputs. This denoising head operates on outputs from the last GrASP block and aims to reduce oversmoothing as over-smoothed outputs can not be used to reconstruct nodes with different features.

#### Postprocessing

The neural network architecture outlined so far scores the likelihood for any given heavy atom to be a part of a binding site as shown in Fig. 3. For applications to drug discovery and model evaluation, it is necessary to aggregate predicted binding site atoms into discrete binding sites. We accomplish this by using average linkage clustering^49^ on all heavy atoms with a predicted binding likelihood above .3. The output clusters are then ranked using the same scoring function as P2Rank except replacing surface points with atoms, 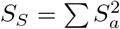where *S*_*S S*_ is the score for a binding site and *S*_*a*_ is the score for an individual atom.^27^ We then obtain the center for each binding site by computing the convex hull of the atom cluster and calculating its center.^54,55^

**FIG. 3:**
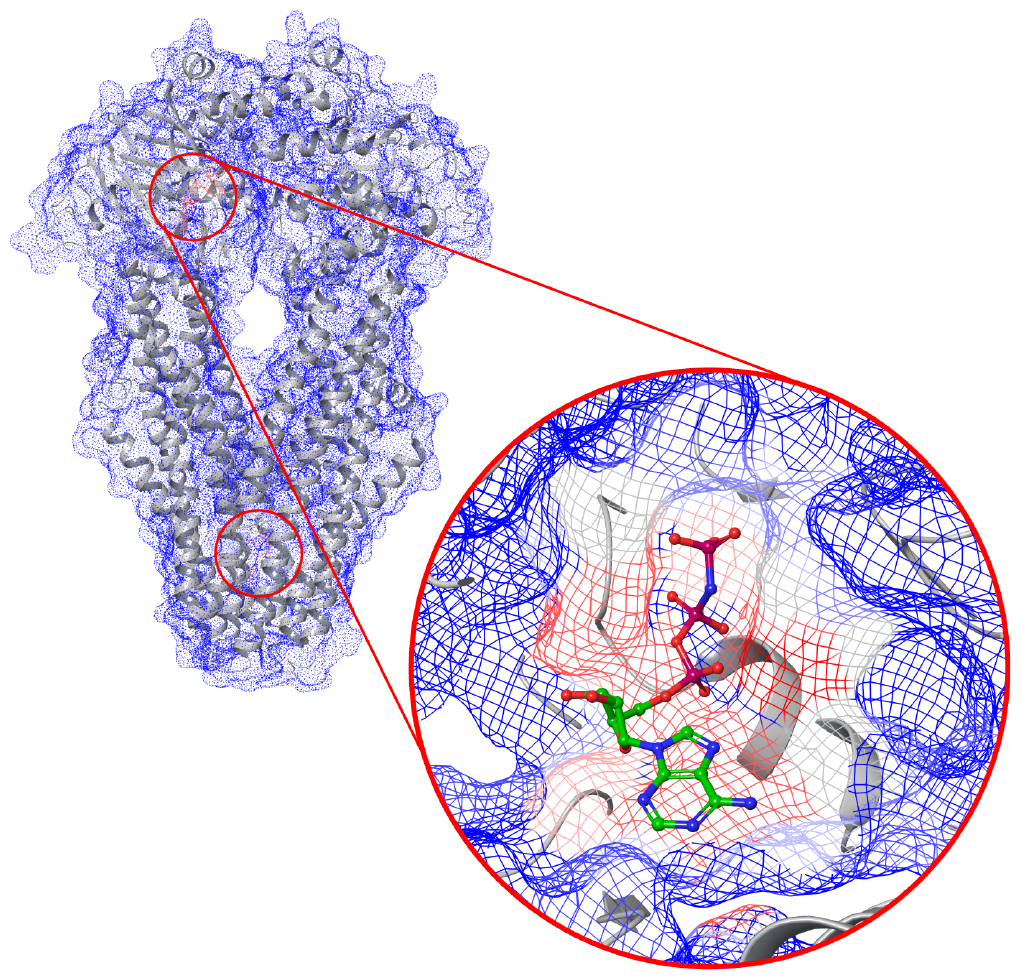
An example of GrASP atom druggability scores ranging from 0 (blue) to 1 (red) for PDB 4Q4A: an ABC transporter that does not have its UniProt ID in GrASP’s training data. High-scoring regions are highlighted with red circles and the scores around the ligand in this structure are shown.

### Relationship to P2Rank

P2Rank is one of the most popular and successful methods for binding site prediction. This method applies a random forest to score points on the protein’s solvent-accessible surface and then aggregates these surface points into sites using single linkage clustering.^27^ While P2Rank uses a different class of model and operates on surface points instead of atoms, P2Rank and GrASP share significant representational similarities. Each surface point in P2Rank describes its local chemical environment using a distance-weighted average of nearby atom properties (up to 6 Å away) with weights 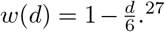 This average can be written as a message passing layer shown in Eq. 4 describing a bipartite graph where surface points *x*_*i*_ receive messages from nearby atoms *x*_*j*_ with distance-based weights shown in Eq. 5. Here we see P2Rank parametrizes the local chemical environment with a single pass through a hand-designed message-passing function. GrASP generalizes this featurization process by learning these aggregation weights through attention and applying multiple message-passing steps.

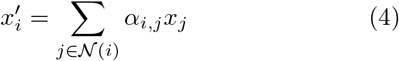

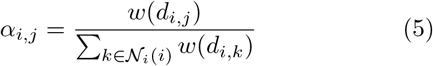

### Datasets

Our training and validation were performed using a modified version of the sc-PDB (v.2017) database.^24^ The sc-PDB is a curated database designed for small ligand docking which contains non-repeating protein-ligand pairs. The crystal structures for these pairs are split into mol2 files which contain the ligand, the binding site (all residues within 6.5 Å), the binding cavity (empty space around the ligand), the full protein, and other structures useful for docking. This database provides 17,594 binding sites and is commonly used to train binding site prediction models but has the shortcoming of unique protein-ligand pairs which means that a large number of binding sites are not labeled. To address this shortcoming, we modify the sc-PDB to contain binding sites corresponding to protein-ligand pairs that are already labeled once (for example, labeling sites on both chains in a symmetric dimer).

We first modify the sc-PDB database by combining entries with the same PDB ID and with protein mol2 files that can be aligned exactly. We then identify unlabeled buried ligands that have the chemical composition as ligands already labeled for any entry with the same PDB ID. We found almost 9,000 additional ligands that fit our criteria which led to a total of 26,196 binding sites across 16,889 protein structures in our final modified dataset. This procedure converts the single-site entries of the sc-PDB into multi-site entries more suitable for binding site prediction methods. The resulting modified dataset is available at github.com/tiwarylab/GrASP and additional details on dataset preparation are available in the SI.

We train and validate our model on the modified dataset with the 10-fold cross-validation splits of the sc-PDB from Ref.^31^ which are made to prevent data leakage with respect to UniProt IDs as well as binding site similarity.

We also modify the test sets used to evaluate P2Rank^27^ to ensure that all ligands are both bound and biologically or pharmacologically relevant. The main preparation of the COACH420 and HOLO4K sets used (i) geometric criteria to ensure the ligand is interacting with the protein, and (ii) simple name filters to avoid the inclusion of water, salt, or sugar as ligands. The P2Rank authors also propose an alternative preparation of these datasets referred to as Mlig sets which use the Binding MOAD database to check that ligands are either biologically or pharmacologically relevant but do not employ previous geometric criteria. We apply both sets of criteria to these sets to ensure both bound and relevant ligands and title the new sets COACH420(Mlig+) and HOLO4K(Mlig+). We also found that HOLO4K contains many multimers with repetitions of the same binding mode. In a real-world setting, multimers would only be considered when they are known to occur *in vivo* and their interface is suspected to be druggable. To reflect this setting, we consider each ligand bound to all proteins within 4 Å and connect all chains that share an interfacial ligand. We then split all systems into subsystems consisting of single chains without interfacial ligands and connected sub-systems with interfacial ligands. This processing should more closely reflect the workflow used in practice avoiding evaluation on homomultimers while preserving evaluation on interfacial binding. The consideration of chains and interfaces does not affect COACH420(Mlig+) as this set only consists of single chains.

## RESULTS

Here we introduce a new metric to evaluate binding site prediction based on standard metrics in semantic segmentation and compare GrASP to P2Rank on updated versions of the original P2Rank datasets.

### Metrics

A commonly used metric to evaluate binding site performance is the distance from the predicted site center to any ligand-heavy atom (DCA). A binding site prediction is considered successful if this distance is below 4 Å and DCA is reported as the percentage of successful predictions over the total number of “ground truth” binding sites (or equivalently bound ligands), usually subject to the constraint that only the top *N* or top *N* + 2 ranked predictions are considered for each system where *N* is the number of binding sites in the ground truth. This metric can be seen as a constrained analogy to recall, a metric commonly used for classification problems defined as 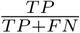 where TP is the number of true positives and FN is the number of false negatives. This ratio can equivalently be defined as the total number of correct predictions divided by the total number of members of the class being predicted. Because DCA refers to both the success criteria and the metric, we will distinguish these two by calling the criteria DCA and the metric DCA recall.

DCA recall evaluates the number of correct predictions among the top *N* binding sites but in a discovery setting the number of binding sites is not known *a priori*. This means that in a real setting any predictions beyond *N* can waste computational resources in downstream tasks even if ranked correctly and likely a fixed maximum number of sites would be considered for each system to stay within a computational budget. To reflect this cost, we propose a constrained analog to the precision metric called DCA precision. DCA precision is the ratio of correctly predicted sites over the total number of predicted sites. This can be computed over all predictions or among the top *M* sites where *M* is a constant that reflects a more realistic cap on the number of sites a user is willing to study per system. DCA precision and DCA recall can be used similarly to the standard precision and recall metrics from machine learning which are always shown together to evaluate the trade-off between false negative and false positive errors.

### Validation Set Results

To evaluate and tune our model we performed 10-fold cross-validation on our augmented sc-PDB database.^31^ The averaged binding site metrics across the 10 folds are shown in Table I with GrASP crossing 90% recall in the top *N* +2 category. Hyperparameter and model architecture choices were made to maximize top *N* DCA recall in this setting.

**TABLE I:** GrASP validation performance averaged across 10 models corresponding to each cross-validation fold in the modified sc-PDB set.

|  | sc-PDB Cross-validation |  |  |  |  |
| --- | --- | --- | --- | --- | --- |
| | DCA Recall<br>Top $N$ (↑) | DCA Recall<br>Top $N + 2$ (↑) | DCA Precision<br>Top 3 (↑) | DCA Precision<br>Top 5 (↑) | DCA Precision<br>All Sites (↑) |
| GrASP | 85.3 | 91.4 | 69.7 | 66.4 | 65.0 |

### Test Set Results

We evaluate both GrASP and P2Rank on our new versions of the COACH420 and HOLO4K sets previously used by P2Rank. COACH420(Mlig+) contains 256 single-chain systems with 315 ligands bound across these systems. This set represents the setting where a small number of predictions are needed and interfacial binding sites are not considered. Table II contains the DCA precision and recall metrics for both methods and shows GrASP has gained 2.6% recall in the top *N* category as well as 30% or greater precision in all categories. To assess the significance of the difference in recall, we used McNemar’s test^56^ comparing which binding sites each method succeeded on. We found that in both the *N* and *N* + 2 categories the difference in recall was not significant. We also assessed the difference in the total number of binding sites returned by each method using the Wilcoxon signed-rank test.^57^ We found the difference in site quantity significant with a p-value less than 0.001 when running three comparisons: comparing the total number of sites, the number in the top 3, and the number in the top 5. This difference explains the contrast in precision between the methods with P2Rank consistently returning more sites. GrASP’s precision is invariant with respect to the number of sites considered in this set while P2Rank’s precision falls as more sites are considered. This difference with respect to the number of sites considered is a consequence of reliance on ranking as there will be many sites returned outside of the top *N*. This shows the necessity of using a maximum number of binding sites and/or a site score threshold when using ranking-based methods in production.

**TABLE II:** Comparison between P2Rank and GrASP performance on the COACH420(Mlig+) test set. Arrows denote whether each metric increases or decreases with higher performance and the highest performance is shown in bold for each metric.

|  | COACH420(Mlig+) |  |  |  |  |
| --- | --- | --- | --- | --- | --- |
| | DCA Recall<br>Top $N$ (↑) | DCA Recall<br>Top $N + 2$ (↑) | DCA Precision<br>Top 3 (↑) | DCA Precision<br>Top 5 (↑) | DCA Precision<br>All Sites (↑) |
| P2Rank | 74.9 | 79.4 | 41.0 | 33.2 | 28.3 |
| GrASP | **77.5** | **80.6** | **71.2** | **71.0** | **71.0** |

HOLO4K(Mlig+) contains a mix of single-chain and multi-chain systems with 6,368 ligands across 4,514 systems. Like COACH420(Mlig+), these systems primarily have one ligand bound, but occasionally contain up to 12 ligands. We show in Table III that GrASP has a similar recall to P2Rank and is even outperformed by 2.2% in top *N* +2 recall but still outperforms P2Rank in precision by a wide margin. We again assessed the significance of these differences using McNemar’s test and the Wilcoxon signed-rank test. The difference in Top *N* recall was not significant but the difference in top *N* + 2 recall was significant with a p-value below 0.001. Similarly, the difference in the number of binding sites was significant with p-value below 0.001 whether considering all sites, the top 3, or the top 5. As before, GrASP’s precision falls by a much smaller amount as more sites are considered, highlighting that ranking too many sites without constraints is insufficient for real-world applications.

**TABLE III:** Comparison between P2Rank and GrASP performance on the HOLO4K(Mlig+) test set. Arrows denote whether each metric increases or decreases with higher performance and the highest performance is shown in bold for each metric.

|  | HOLO4K(Mlig+) |  |  |  |  |
| --- | --- | --- | --- | --- | --- |
| | DCA Recall<br>Top $N$ (↑) | DCA Recall<br>Top $N + 2$ (↑) | DCA Precision<br>Top 3 (↑) | DCA Precision<br>Top 5 (↑) | DCA Precision<br>All Sites (↑) |
| P2Rank | 81.2 | **86.5** | 45.9 | 35.4 | 25.5 |
| GrASP | **81.3** | 84.3 | **72.8** | **71.6** | **71.4** |

While computing the contingency tables for McNemar’s test, we saw that many of the binding sites that were failure cases for one method were successes for the other. This prompted us to calculate the percentage of binding sites where either GrASP or P2Rank are successful. For *N* and *N* + 2 on COACH420(Mlig+) either method succeeded on 84.13% and 86.35 % of sites respectively. For HOLO4K(Mlig+) either method succeeded on 90.33% for top *N* and 92.73% for top *N* + 2. Using predictions from both models provides a significant increase in binding site coverage and may be beneficial in studies where precision isn’t valued.

We also compute DCA recall with varying success thresholds for both test sets in Fig. 4. Interestingly with less strict DCA success thresholds, P2Rank outperforms GrASP on both top *N* and *N* + 2 on HOLO4K(Mlig+) but GrASP’s top *N* recall improves so significantly COACH420(Mlig+) that it outperforms P2Rank’s top *N* + 2 recall.

**FIG. 4:**
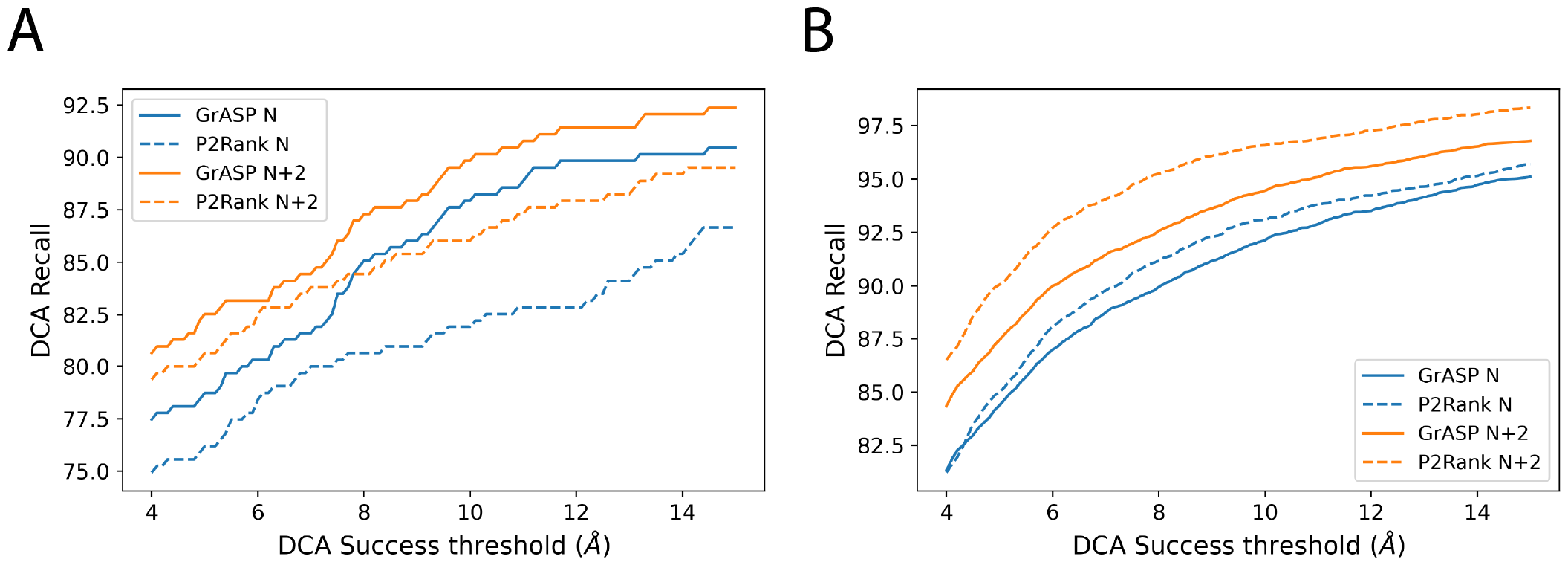
Comparison of DCA recall for GrASP and P2Rank with varying DCA success thresholds for A) COACH420(Mlig+) and B) HOLO4K(Mlig+).

### Sequence Identity Generalization

The UniProt splitting criterion commonly used to prevent leakage between train and test sets is insufficient to assess a model’s ability to generalize to novel proteins. While this approach mirrors the original P2Rank approach, we can quantify generalization more carefully by analyzing success rates as a function of sequence identity between the train and test sets. We used MMseqs2^58^ to find the most similar entry in the training set for each system in the test set and assigned this sequence identity to all labeled binding sites in the test system. We then assigned each test binding site into histogram bins with 10% intervals in sequence identity (including the lower bound but not the upper bound). We recalculated top *N* and *N* + 2 DCA recall for each sequence identity bin individually to assess GrASP’s performance with respect to the novelty of the test system’s sequence. We show in Fig. 5 that GrASP’s DCA recall has a very small variance with respect to sequence identify for all bins with sufficient data (above 20% identity). Notably, GrASP is still able to maintain the same success rate for the 20-30% range where proteins are much less likely to be homologous. Here we show this analysis for GrASP on HOLO4K(Mlig+) because the size of the test set allows for small standard error but we show this analysis for both GrASP and P2Rank on both test sets in the SI. P2Rank’s performance is also similar in all well-sampled bins but with higher variance than GrASP.

**FIG. 5:**
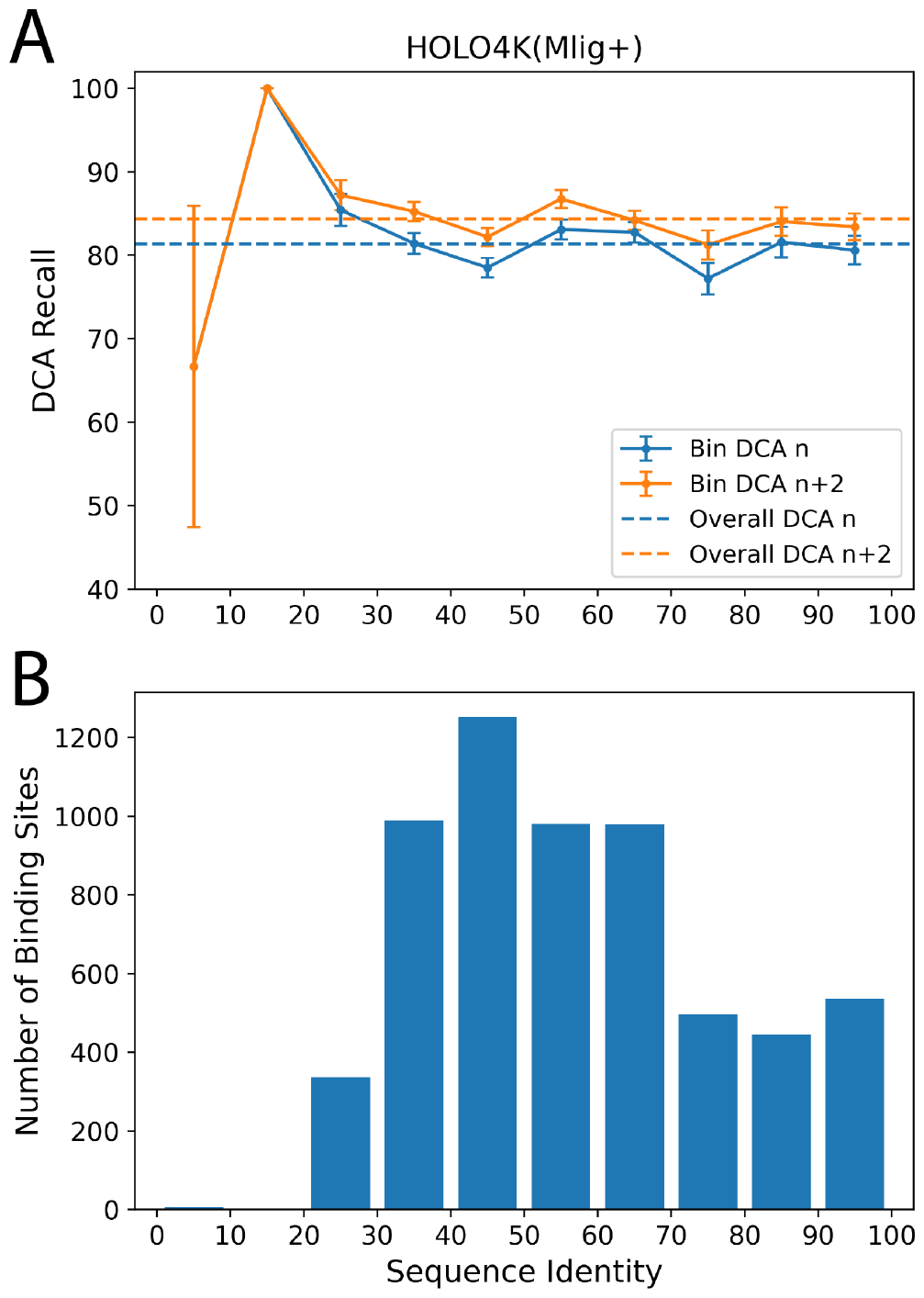
GrASP’s performance on the HOLO4K(Mlig+) set as a function of sequence similarity between train and test sets. A) GrASP’s performance on samples in each sequence similarity bin with standard error is displayed as bars and the performance on the full set is shown as dashed lines. B) Histogram of sequence similarity between GrASP’s training data and HOLO4K(Mlig+). Note that the 0-20% range has insufficient data to draw meaningful conclusions.

## DISCUSSION

In this work, we have developed a new method called Graph Attention Site Prediction which reaches state-of-the-art performance in binding site recall and does so with much higher high precision, a metric that has not yet been reported for binding site prediction, but affects the computational cost to use predicted binding sites for other tasks. Precision analysis in the setting where the number of binding sites is unknown shows a weakness of ranking-based methods. If the true number of sites is not known there is not a clear stopping point when using a ranked list and downstream tasks may be frequently performed on poor predictions. We predict that coupling a ranked binding site list with a site score threshold to discard poor predictions would improve precision, and in turn, reduce waste in downstream tasks for drug discovery. We recommend future methods aim to optimize such thresholds and report both precision and recall for DCA or other metrics of their choice.

Currently, binding site prediction methods either rank binding sites generated with geometric criteria or perform semantic segmentation and then cluster the segmentation mask. Future methods should treat binding site prediction as an instance segmentation task where the model predicts both which atoms (or surface points) are part of a binding site and which binding site they belong to. The current clustering-based instance segmentation is not end-to-end differentiable and lags behind the methodology used in image segmentation.^59^ Given this suboptimal step in current methods, we recommend that small-scale projects use the raw semantic segmentation scores on surface atoms and hand-pick where to dock ligands. We also recommend that the community increases focus on treating the task as instance segmentation instead of perfecting methods for semantic segmentation because clustering quality may set a cap on performance.

## DATA AVAILABILITY STATEMENT

The trained GrASP model together with code, an easy-to-use web interface through Google Colab, and associated datasets to retrain the model are available at github.com/tiwarylab/GrASP.

## ACKNOWLEDGMENTS

The research reported in this publication was supported by the National Institute Of General Medical Sciences of the National Institutes of Health under Award Number R35GM142719. The content is solely the responsibility of the authors and does not necessarily represent the official views of the National Institutes of Health. We are grateful to NSF ACCESS Bridges2 (project CHE180053) and University of Maryland Zaratan High-Performance Computing cluster for enabling the work performed here. The authors thank the P2Rank team for discussing their data sets, thank Michelle Girvan for suggesting average linkage clustering, thank Schrödinger and Nimbus Therapeutics scientists for discussing test set preparation, and thank Pavan Ravindra for discussing GNNs.

## NOTES

The authors declare the following competing financial interest(s): P.T. is a consultant to Schrödinger, Inc. and is on their Scientific Advisory Board.

## SUPPORTING INFORMATION AVAILABLE

Supporting Information contains a detailed description of methods and additional model evaluation.

---

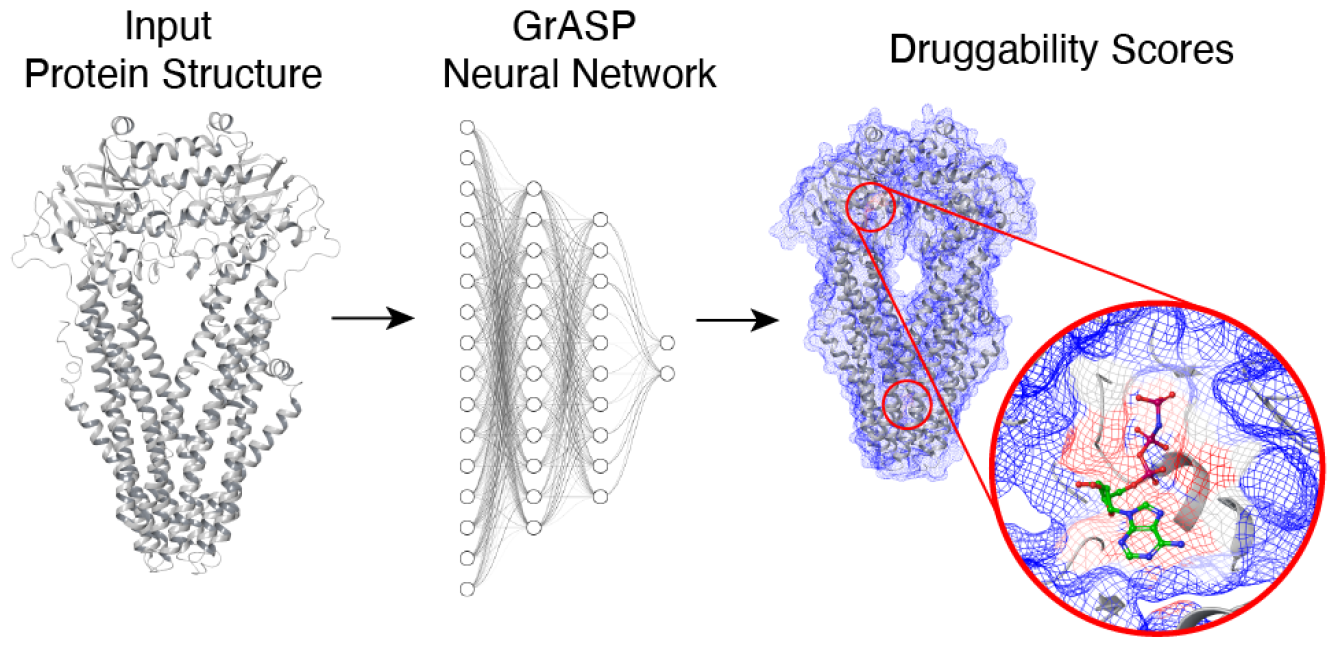

## Notes

### Competing Interest Statement

P.T. is a consultant to Schrodinger, Inc. and is on their Scientific Advisory Board.

### Summary of Updates

Added one new figure and further technical details

