## Supplementary Materials for "Graph Attention Site Prediction (GrASP): Identifying Druggable Binding Sites Using Graph Neural Networks with Attention"

| **Atom-Scale Features** |
| --- |
| Atom Type |
| Local Density of Atoms (9 features, density within spheres ranging from 2 - 10 Å) |
| Solvent Accessible Surface Area |
| Formal Charge |
| Number of Bonds with Heavy Atom |
| Ring Membership |
| Aromaticity |
| Mass |
| Hybridization |
| Hydrogen Bond Donor/Acceptor |
| Hydrophobicity |
| **Amino Acid-Scale Features** |
| Residue Name |
| Residue Polarity |
| Acidity (Acidic/Basic/Neutral) |
| Charge (Positive/Negative/Neutral) |
| **Edge Features** |
| Inverse Distance |
| Bond Order (including unbonded) |

**TABLE S1:** GrASP input features.

### I. SC-PDB DATASET PREPARATION

Unlabeled ligands in the modified sc-PDB dataset were identified by matching their chemical composition to labeled ligands from an entry with the same PDB ID. This criterion was chosen to avoid adding ligands not present in the original sc-PDB to be consistent with the sc-PDB's requirement that ligands must be biologically relevant. The count of each non-hydrogen element in the ligand was compared to the labeled ligands and matches were recorded. The relatively loose criteria of element count was chosen to avoid false negatives due to inconsistencies in bond perception and ligands with a different ordering of elements, different sybyl atom types, or different residue names were labeled for visual inspection to confirm they were not identified as duplicates in error.

Unlabeled ligands were identified as buried using the ratio of solvent-accessible surface area (SASA) in the protein complex to SASA in vacuum. Ligands were classified as buried if they were either below 30% solvent accessible or if their fraction accessible was no more than 10% above their labeled counterpart's fraction. These criteria use a conservative definition of buried ligands with 30% accessible surface chosen because 95% of the labeled ligands fall below this threshold. The second part of the criteria accounts for ligands that have over 30% surface exposure in their binding modes. We found that ligands with long tails may be up to 60% accessible in their labeled binding mode and this comparison to the labeled ligands identifies the small number of cases where unlabeled ligands are symmetric to their labeled counterparts but over 30% accessible. The use of two criteria allows the base threshold of 30% to be low enough to avoid false positives while comparison to the labeled fraction accessible catches false negatives that would arise due to unique binding modes.

### II. MOTIVATION FOR DATASET CHANGES

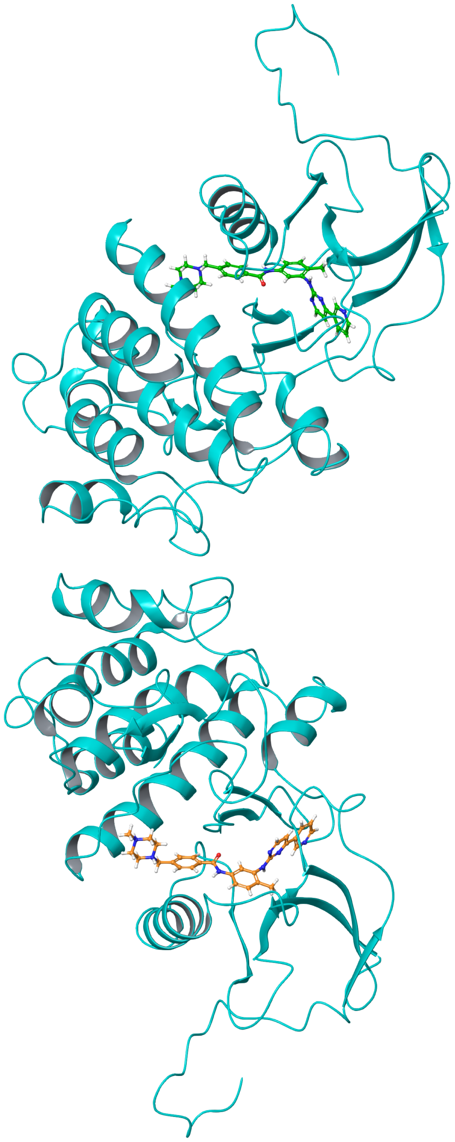

**FIG. S1:** Example of the motivation for parsing new ligands in the sc-PDB dataset (PDB: 1IEP). The ligand mol2 file in the original dataset contains the green ligand while the protein file contains both the cyan protein and orange ligand. Naive use of the dataset may lead a modeler to label no binding site in the lower chain and potentially include the ligand in the receptor inputs. Our new preparation will return two ligand structure files separating these ligands and a protein file containing the cyan receptor.

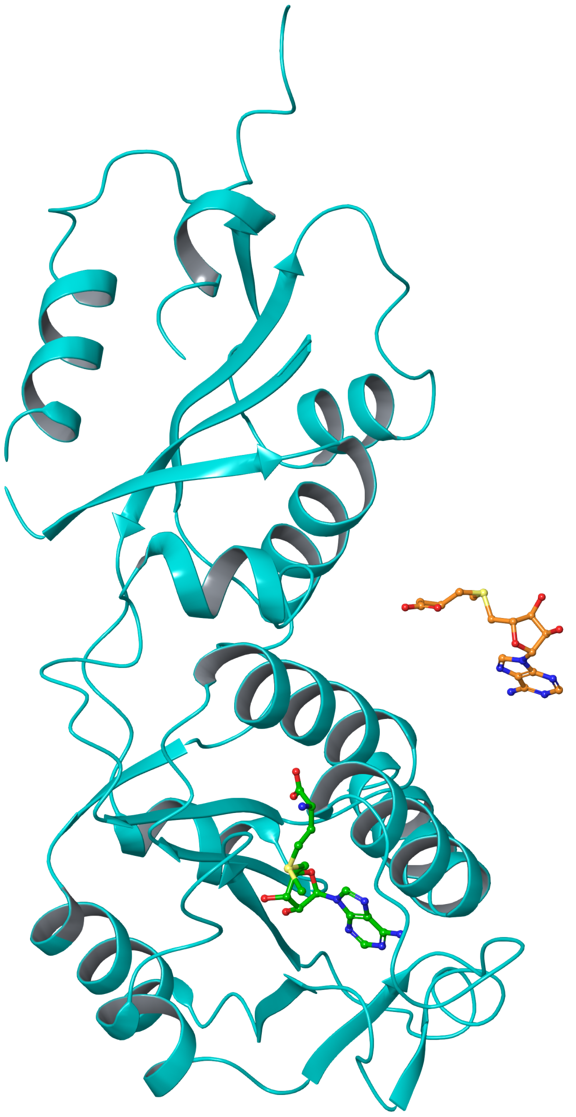

**FIG. S2:** Example of the motivation to combine the biological relevance and geometric criteria used separately by P2Rank (PDB: 3NK7 in COACH420). This structure was prepared using only the biological relevance criteria that the ligand must be present in the binding MOAD database.[1] The orange ligand does not meet the geometric criteria to consider it bound and would only be included in one of the ways that P2Rank parses COACH420 (the Mlig version). To avoid such cases while guaranteeing the relevance of the ligands, we only include ligands, such as the green ligand, that pass both sets of criteria. Practically this is accomplished by parsing each test set with P2Rank once with each type of criteria and then taking the ligands included in both preparations.

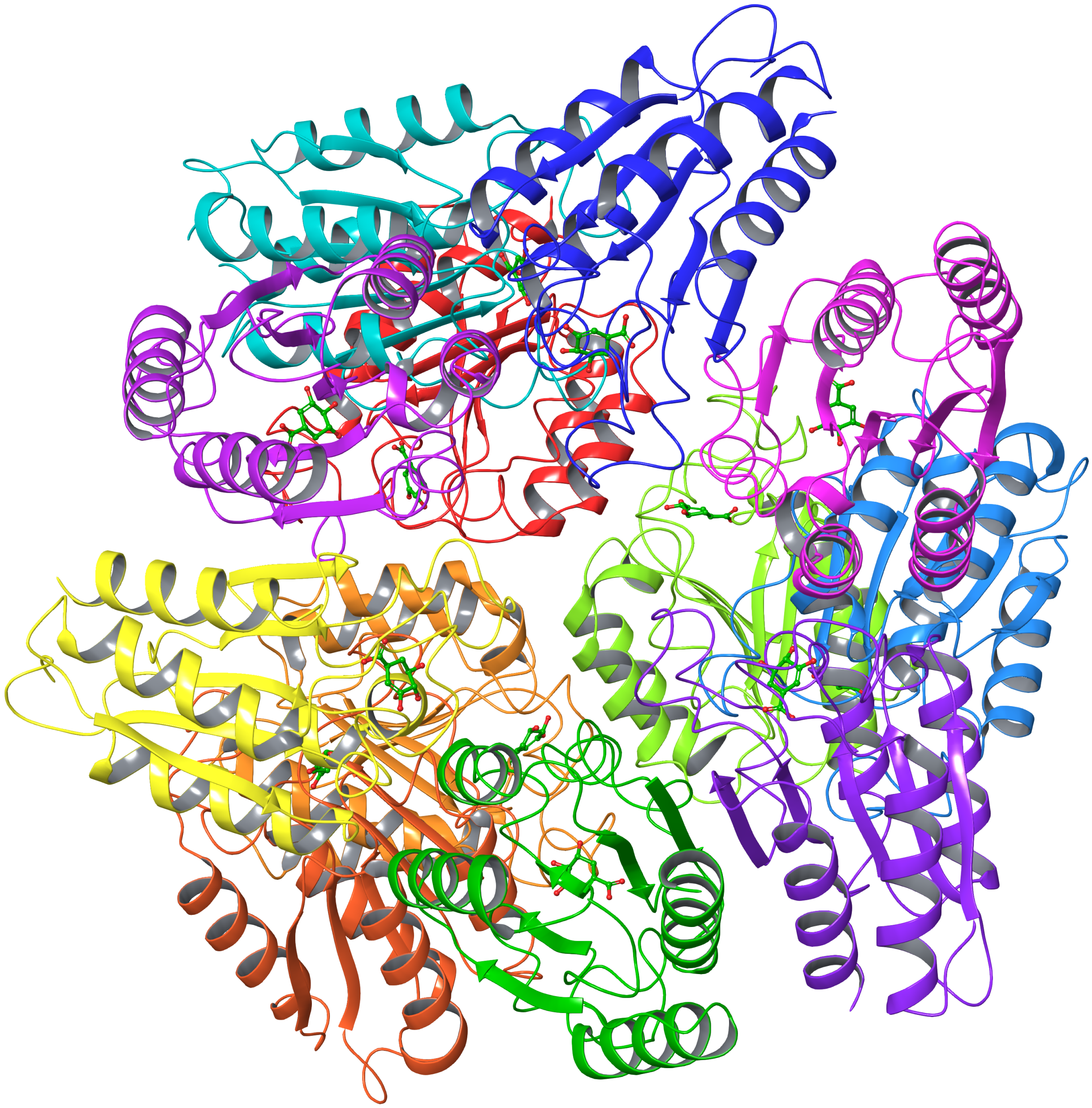

**FIG. S3:** Example of the motivation for our methodology to split chains in the test sets. This system (PDB: 1GTZ) in the HOLO4K set is a homo-dodecamer with each of the 12 chains shown in a different color. Since it is unlikely that all 12 chains would be used at once in a virtual screen, it is more realistic to evaluate predictions on a subsystem. Naively we could make predictions on each chain individually but this would misrepresent binding sites at the interface between chains. We therefore separate chains that do have a ligand at their interface. Specifically, we associate each ligand with all chains within 4 Å (the threshold from P2Rank's geometric binding criteria) to determine which ligands bind to multiple chains.

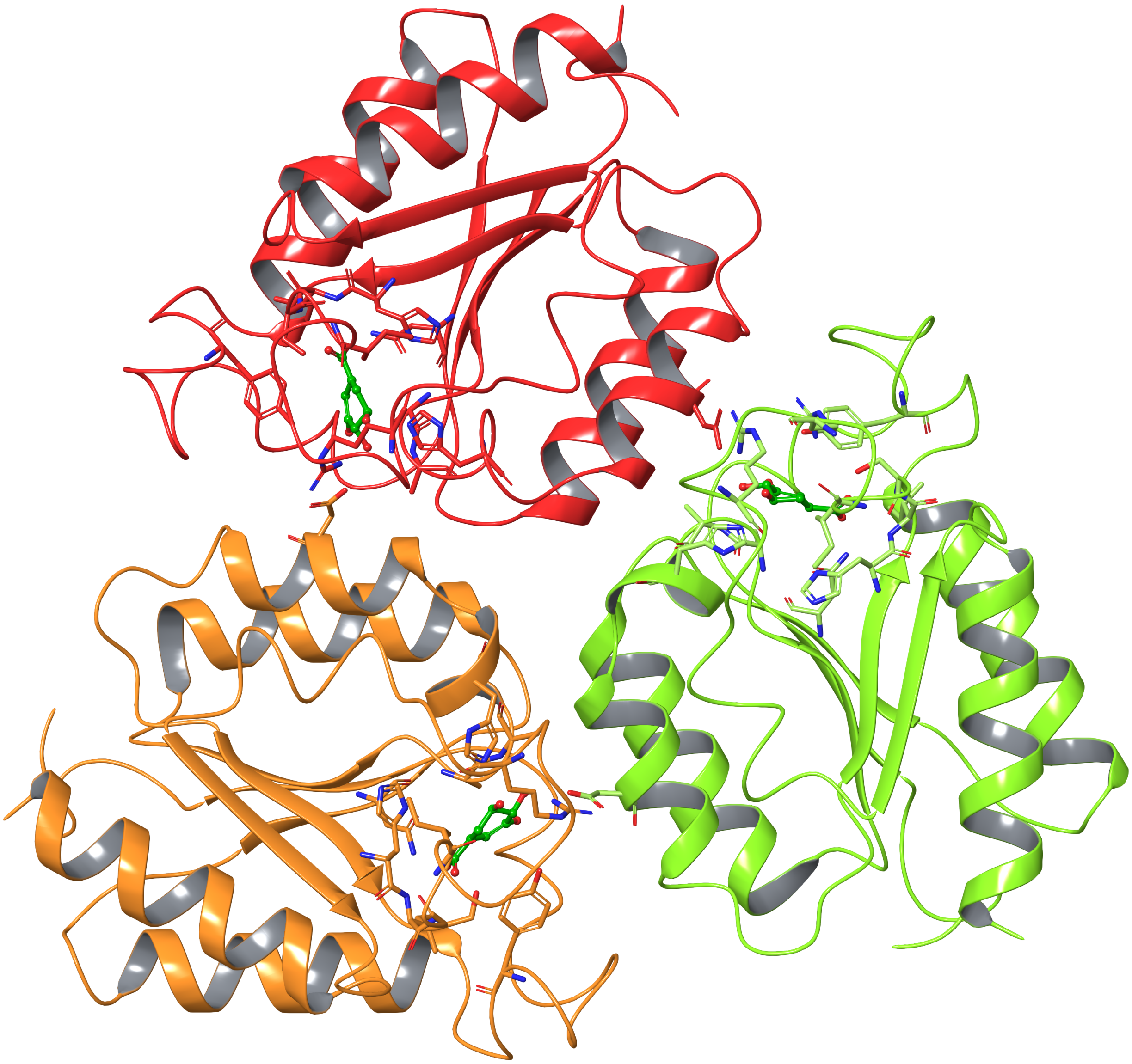

**FIG. S4:** A subsystem created from the system (PDB: 1GTZ) in Fig. S3 above. We color the protein by chain and show all protein residues within 4 Å of the ligands. We can see that each ligand is close to two chains so we do not separate any of these chain pairs. Overall, our parsing splits the homo-dodecamer into four subsystems with three chains and three ligands each. We choose to include all subsystems after splitting chains so this change does not affect the distribution of ligands in the test set but penalizes ranking multimer binding sites in a more realistic setting.

### III. NEAR-SURFACE GRAPH CONSTRUCTION

The near-surface graph is constructed using two components: a distance matrix between all protein heavy atoms, and labels for which heavy atoms are part of the protein surface. We begin the process of marking which atoms are on the surface by first calculating the SASA. We then add the hydrogen SASA to their bonded heavy atoms since GrASP will use implicit hydrogens. We apply a threshold of $10^{-4}$ $nm^2$, which was treated as a hyperparameter, marking all atoms with SASA above this threshold as part of the surface. Once we have calculated a distance matrix and have labelled the surface atoms, we take the set of all surface heavy-atoms and add them as nodes to the protein graph. We then add all heavy-atoms within 5 Å of surface heavy-atoms as nodes. Finally, we draw edges between all nodes corresponding to atoms $\leq$ 5 Å apart.

### IV. BINDING SITE LABELS

We train GrASP using continuously valued binding site labels with the following sigmoid form where $y_i$ is the class label for protein atom $i$ and $d_i$ is the distance from protein atom $i$ to the nearest ligand heavy atom. Only surface atoms defined in Section III are scored by GrASP.

$$y_i = \text{Sigmoid}(-3(d_i - 5)) \tag{1}$$

This can be viewed as a smoothed version of a 5 Å binding site definition where the labels decrease from 1 to 0 in roughly the region between 4 - 6 Å as opposed to a discrete boundary at 5 Å. Both the midpoint and slope of this sigmoid were tuned as hyperparameters to optimize the top $N$ DCA recall on the validation set.

### V. CONVEX HULL CENTER CALCULATION

To calculate the center of a convex hull, the hull is treated as a solid object with uniform density and its center of mass is calculated. This is accomplished by breaking the hull into tetrahedrons and taking the volume-weighted average of these tetrahedrons' centroids.

### VI. SEMANTIC SEGMENTATION METRICS

We use the following metrics to evaluate the performance of semantic segmentation:

- Area under the receiver operating characteristic curve (ROC AUC): A classification threshold invariant metric that measures the trade-off between the true positive rate and false positive rate as the classification threshold is varied. Here we use macro averaging to give equal weight to each category instead of each sample because there is heavy class imbalance between site and non-site atoms.
- Area under the precision-recall curve (PR AUC): A classification threshold invariant metric that measures the trade-off between precision and recall as the classification threshold is varied. Similar to ROC AUC but less sensitive to class imbalance.
- Matthews correlation coefficient[2] (MCC): A correlation metric commonly used for tasks where there is class imbalance. A value of 1 means all predictions are correct while a value of -1 means all predictions are incorrect and a value of 0 corresponds to random predictions. The MCC is shown in Eq. 2 where T, F, P, and N correspond to true, false, positive, and negative respectively.

$$MCC = \frac{TP \times TN - FP \times FN}{\sqrt{(TP + FP)(TP + FN)(TN + FP)(TN + FN)}} \tag{2}$$

| COACH420(Mlig+) | | | HOLO4K(Mlig+) | | |
|---|---|---|---|---|---|
| ROC AUC | PR AUC | MCC | ROC AUC | PR AUC | MCC |
| .97 | .75 | .64 | .98 | .79 | .70 |

**TABLE S2:** GrASP atom-wise semantic segmentation metrics on COACH420(Mlig+) and HOLO4K(Mlig+).

### VII. SEQUENCE IDENTITY GENERALIZATION

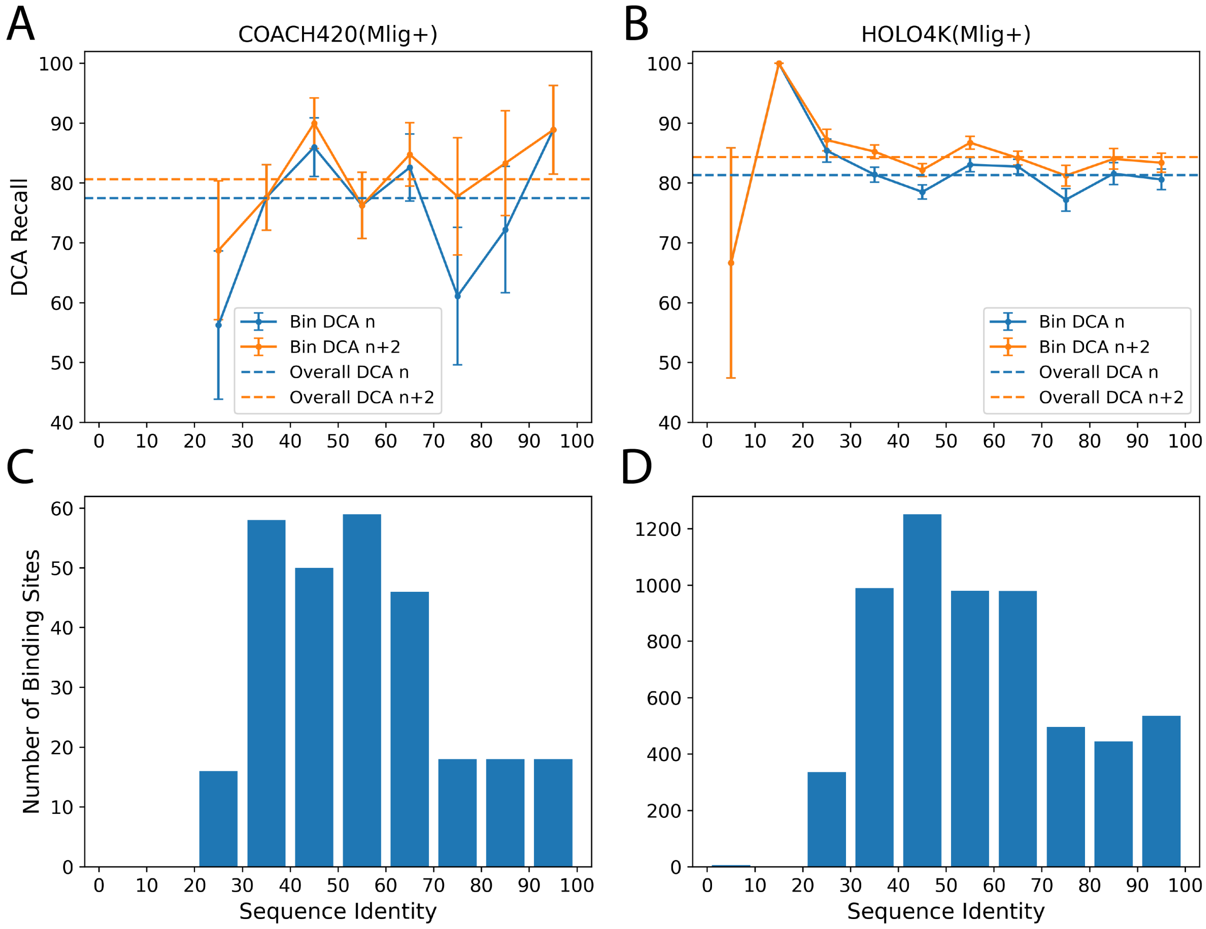


**FIG. S5:** GrASP's performance on the COACH420(Mlig+) and HOLO4K(Mlig+) sets as a function of sequence similarity between train and test sets. GrASP's performance on samples in each sequence similarity bin with standard error is displayed as bars and the performance on the full set is shown as dashed lines for A) COACH420(Mlig+) and B) HOLO4K(Mlig+). Histogram of sequence similarity between GrASP's training data and C) COACH420(Mlig+) or D) HOLO4K(Mlig+).

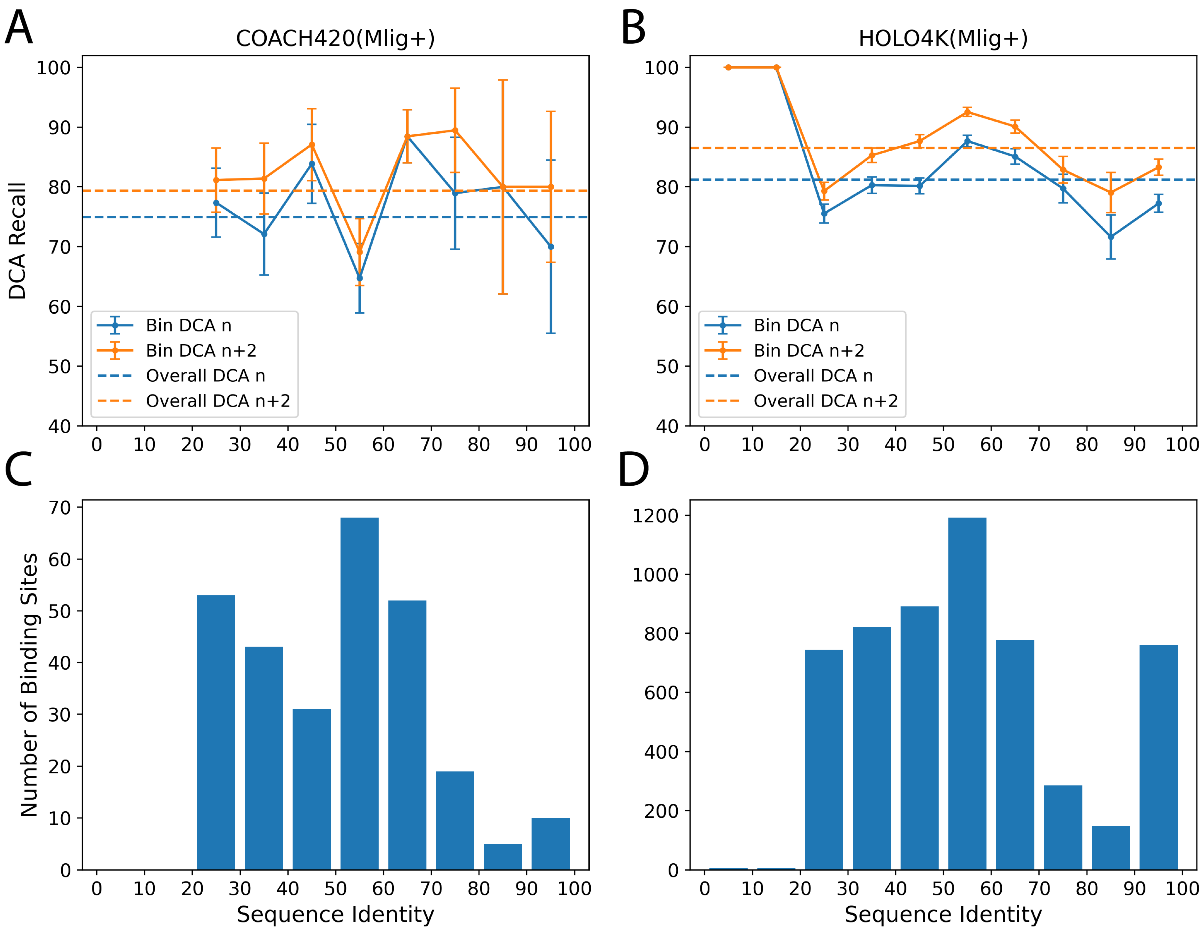


**FIG. S6:** P2Rank's performance on the COACH420(Mlig+) and HOLO4K(Mlig+) sets as a function of sequence similarity between train and test sets. P2Rank's performance on samples in each sequence similarity bin with standard error is displayed as bars and the performance on the full set is shown as dashed lines for A) COACH420(Mlig+) and B) HOLO4K(Mlig+). Histogram of sequence similarity between P2Rank's training data and C) COACH420(Mlig+) or D) HOLO4K(Mlig+).

### VIII. PREDICTIONS ON APO STRUCTURES

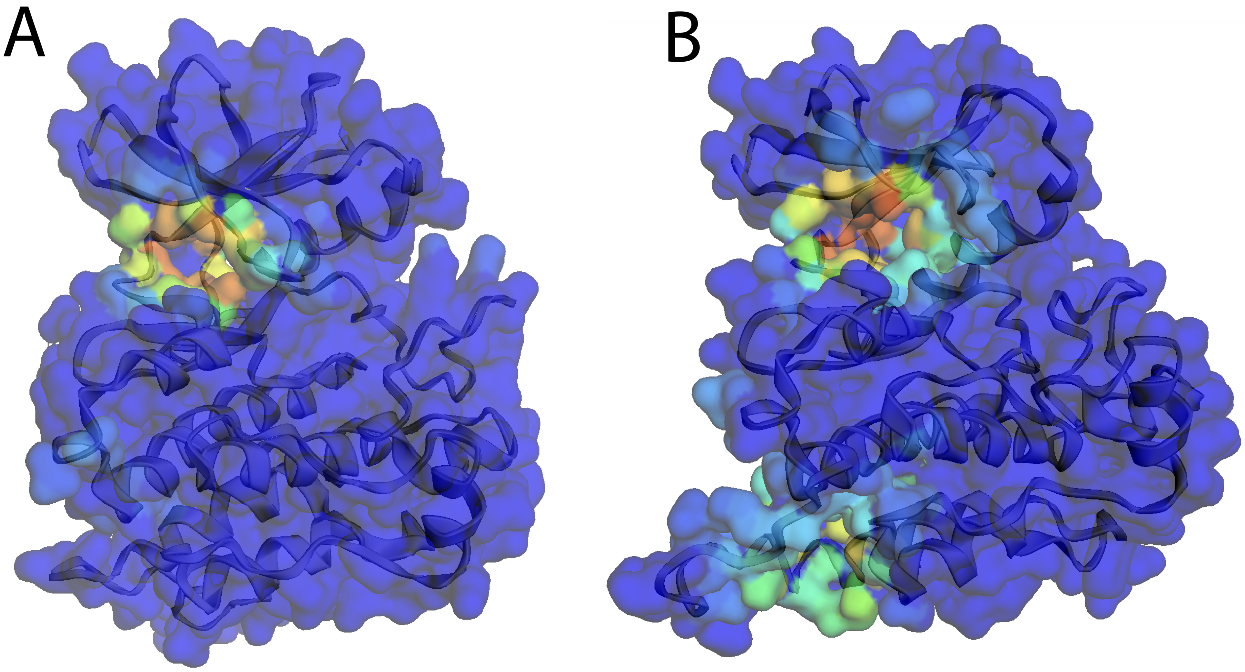

**FIG. S7:** Example outputs of the GrASP web interface on two apo structures of Abl Kinase in the DFG-in (A. PDB: 6XR6) and DFG-out (B. PDB: 6XR7) conformations. The druggability scores for each surface atom are used to color the transparent surface with dark blue corresponding to a score of 0 and dark orange corresponding to a score of 1. While Abl Kinase is included in the training set, no apo structures were included in the training set. These preliminary results show that GrASP still identifies the accessible portion of the orthosteric binding site in both DFG-in and DFG-out apo conformations despite the site being more occluded than in training. It also appears that GrASP identifies the allosteric site in the DFG-out conformation with lower confidence than the orthosteric site. These initial results show some promise for GrASP's potential to identify binding sites on holo and apo structures in a conformation-dependent manner. This potential could be used for docking structures where no site is known or conformations where a cryptic site may be exposed. These settings will be explored in a rigorous quantitative manner in a future publication.

### IX. ABLATION STUDIES

We assessed the contribution of individual GrASP components by performing ablation studies removing one or more components. The ablation models we used were as follows:

- No Noisy Nodes: GrASP trained without the Noisy Nodes reconstruction loss.
- Sum Aggr: GrASP without multi-aggregation using sum aggregation like the original GAT.
- Mean Aggr: GrASP without multi-aggregation using mean aggregation.
- No Skip Connections: GrASP without residual or jumping knowledge skip connections.
- GATv1 GrASP: GrASP using the original GAT instead of GATv2.
- GCN GrASP: GrASP using GCN instead of GATv2.
- Simplified GrASP Sum Aggr: GrASP without skip connections, noisy nodes or multi-aggregation using sum aggregation.
- Simplified GrASP Mean Aggr: GrASP without skip connections, noisy nodes or multi-aggregation using mean aggregation.
- No Dropout: GrASP without dropout in the attention weights.
- Batch Norm: GrASP using batch norm instead of instance norm.
- No Norm: GrASP with instance norm removed.
- *Discrete Sites: GrASP with discrete site definition instead of continuous.
- *Full Graph Inference: GrASP trained to score all atoms in the graph without considering solvent accessibility.

We trained these ablation models on a single validation fold as well as the training splits associated with COACH420(Mlig+) and HOLO4K(Mlig+). We then assessed their performance at identifying which atoms were part of binding sites using MCC and PR-AUC. Note that we exclude the two ablation models marked with an asterisk, discrete sites and full graph inference, from atom scoring comparisons because they change the definition of binding site used to train the model and do not allow for fair comparison on the atomic scale. We then cluster atomic scores into binding sites and compare the DCA precision and recall for each model and set. We obtain p-values comparing default GrASP to each model for atom metrics using McNemar's test[3] on their atom labels. We obtain p-values for DCA recall using McNemar's test on the binding site predictions and we obtain p-values for the number of sites predicted (shown above precision for convenience) using the Wilcoxon signed-rank test.[4] All p-values were then corrected using the Holm–Bonferroni method[5] to control the family-wise error rate for our set of models and associated hypotheses.

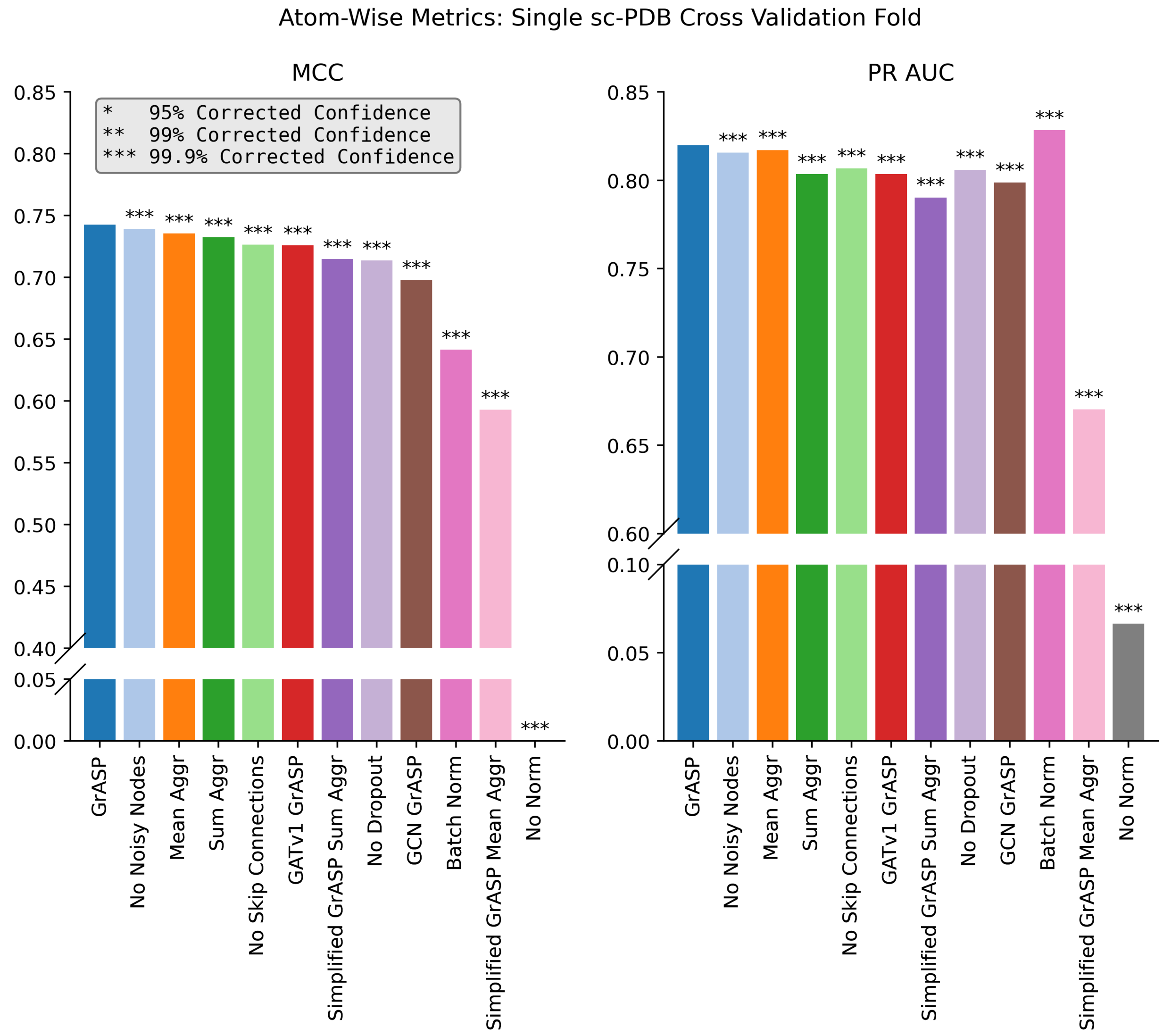


**FIG. S8:** Atom metrics for a single sc-PDB cross-validation fold. Models are ordered based on their MCC performance and confidence in the difference between default GrASP and each model is shown with asterisks. We can see that GrASP significantly outperforms all ablation models in MCC and is only outperformed by the Batch Norm model in PR AUC.

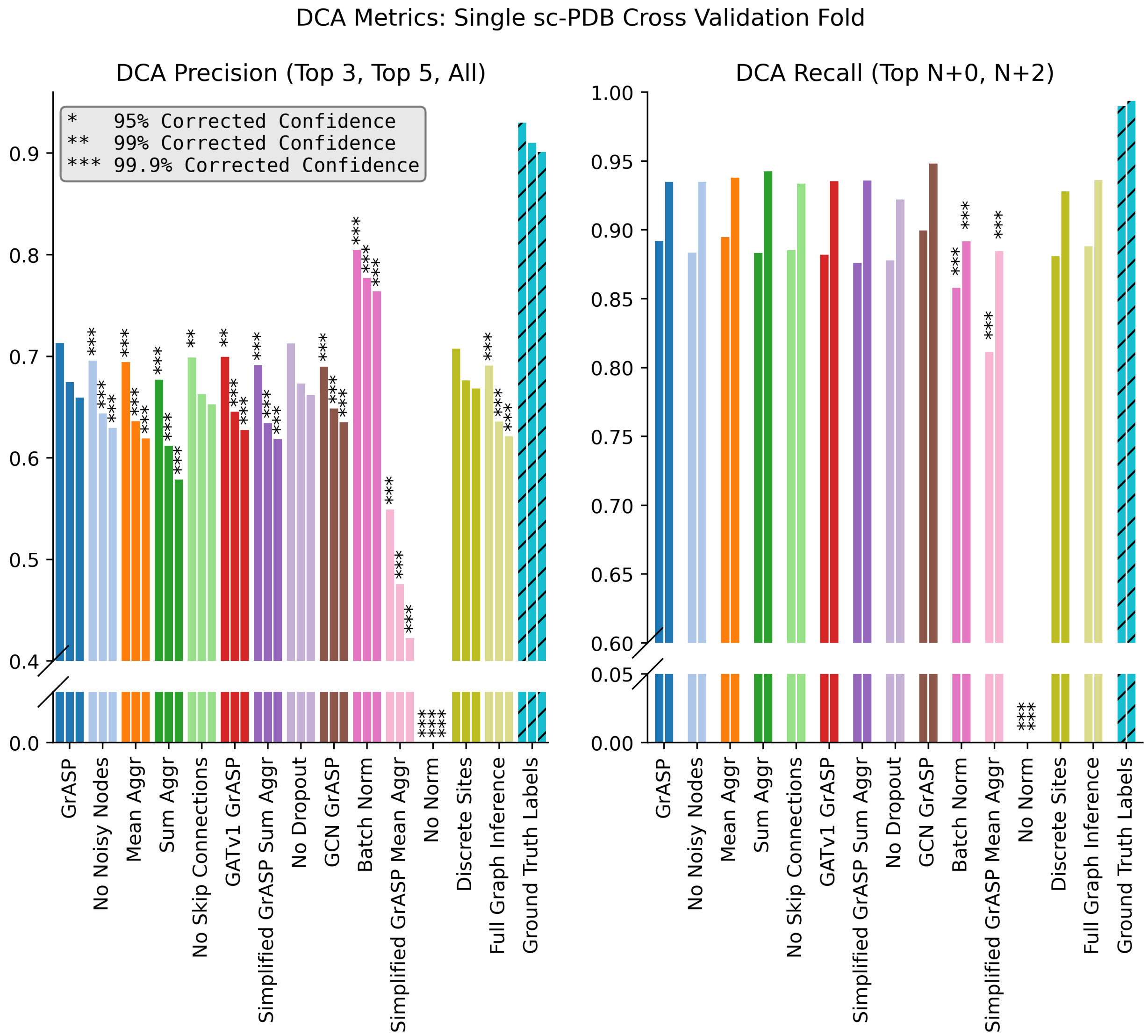


**FIG. S9:** DCA metrics for a single sc-PDB cross-validation fold with clustering performance on training labels shown as striped bars for reference. Models are ordered based on their validation MCC performance and confidence in the difference between default GrASP and each model is shown with asterisks. The asterisks above precision represent differences in the number of sites predicted. We can see that no model significantly outperforms GrASP in DCA recall and that the increased PR AUC previously noted for Batch Norm corresponds to a precision-recall trade-off. It is important to note that significant improvements in atom scoring over the ablation models do not translate to significant improvements in DCA.

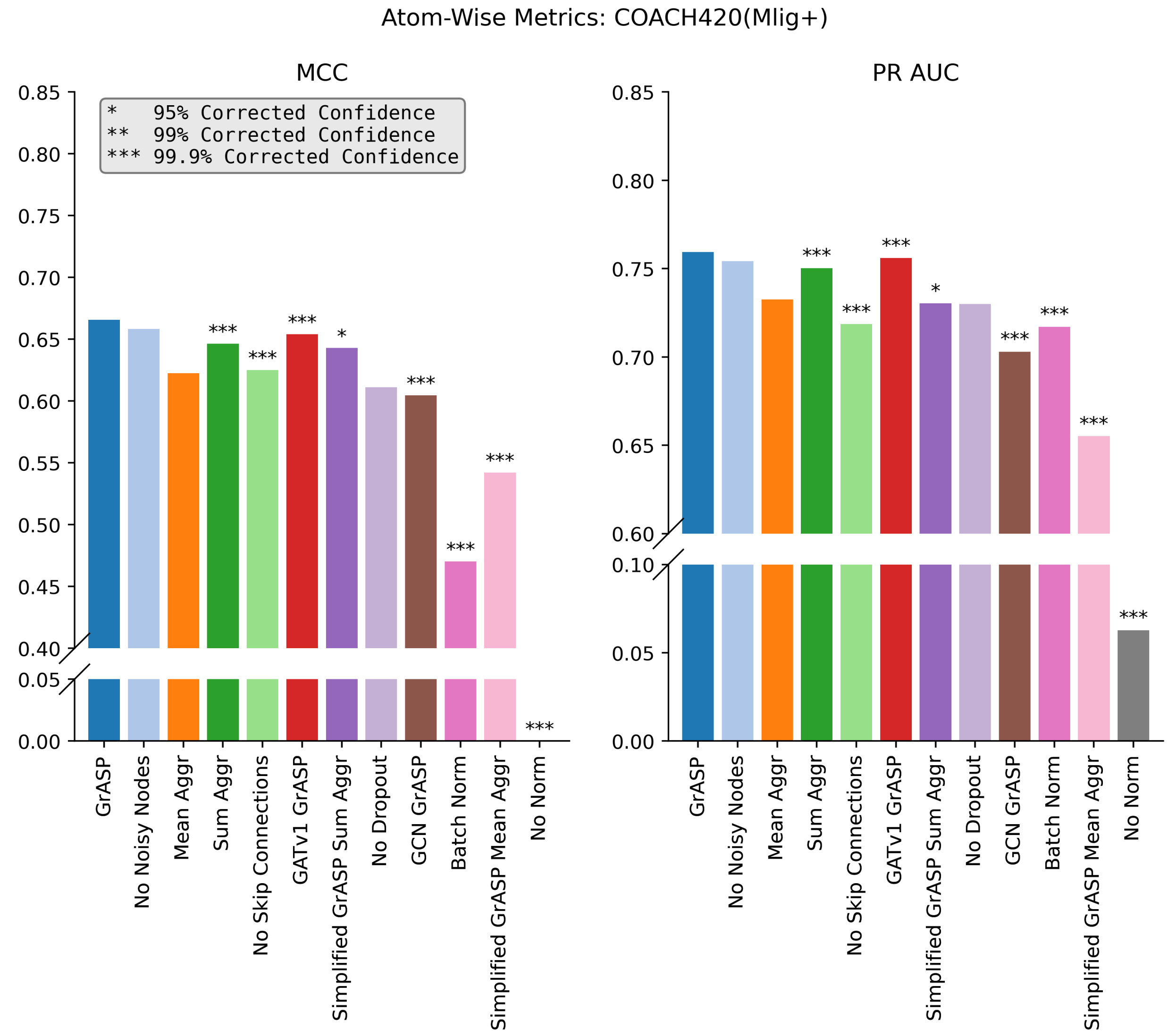


**FIG. S10:** Atom metrics for COACH420(Mlig+). Models are ordered based on their validation MCC performance and confidence in the difference between default GrASP and each model is shown with asterisks. We can see that no model outperforms the original GrASP but with the smaller data set the differences are not significant for some differences.

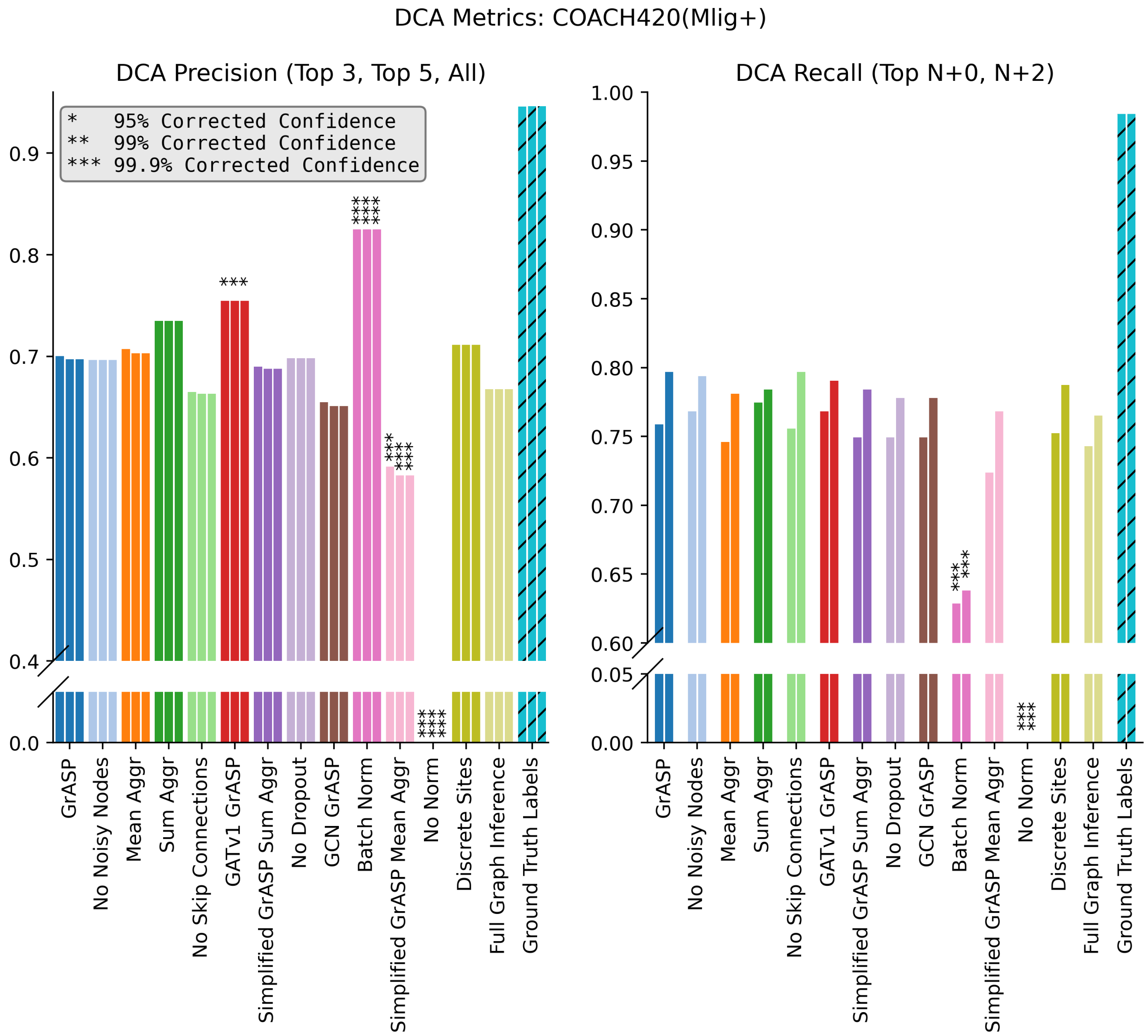


**FIG. S11:** DCA metrics for COACH420(Mlig+) with clustering performance on training labels shown as striped bars for reference. Models are ordered based on their validation MCC performance and confidence in the difference between default GrASP and each model is shown with asterisks. The asterisks above precision represent differences in the number of sites predicted. While default GrASP has the highest DCA recall it is not significantly higher than most ablation models and GrASP is significantly outperformed by GATv1 GrASP and Batch Norm in precision. As before, we note that significant improvements in atom scoring over the ablation models do not translate to significant improvements in DCA.

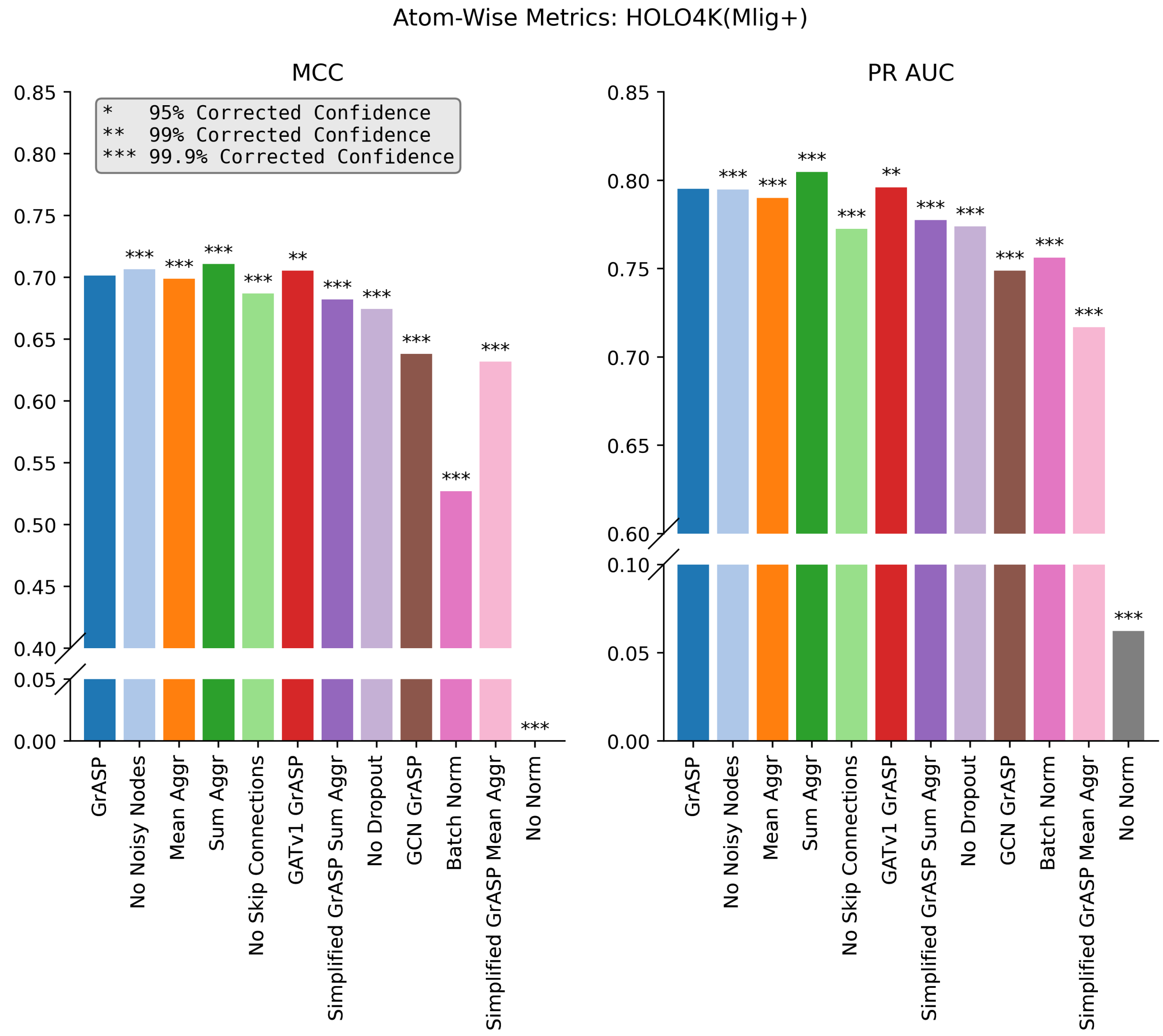


**FIG. S12:** Atom metrics for HOLO4K(Mlig+). Models are ordered based on their validation MCC performance and confidence in the difference between default GrASP and each model is shown with asterisks. We can see that GrASP significantly outperforms most models but on this dataset No Noisy Nodes, Sum Aggr, and Simplified GrASP Sum Aggr outperform the original model.

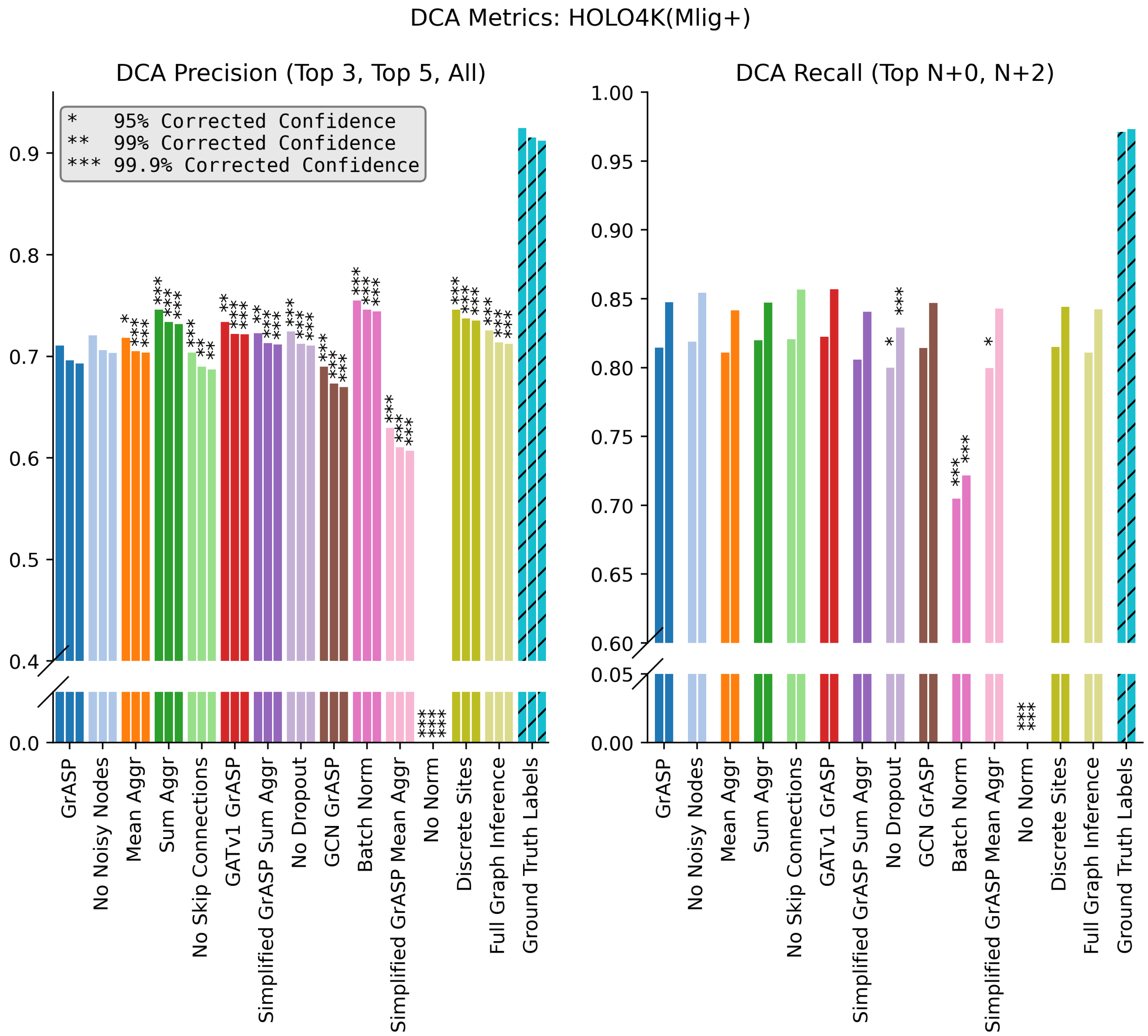


**FIG. S13:** DCA metrics for HOLO4K(Mlig+) with clustering performance on training labels shown as striped bars for reference. Models are ordered based on their validation MCC performance and confidence in the difference between default GrASP and each model is shown with asterisks. The asterisks above precision represent differences in the number of sites predicted. GrASP is not significantly outperformed by any models in DCA recall, however, several models have significantly higher precision. As before, we note that significant differences in atom scoring over the ablation models do not translate to significant differences in DCA.
